# Molecular characterization and pathogenicity assessment of bacteria causing infectious diseases in cage-farmed fish in the Lake Victoria

**DOI:** 10.1101/2025.03.02.641053

**Authors:** Magoti Ernest Ndaro, Jackson L. Saiperaki, Alexanda Mzula, Silvia Materu, Antony Funga, Beda J. Mwang’onde, Catherine A. Mwakosya, Cyrus Rumisha

## Abstract

Despite government initiatives in East African countries promoting cage fish farming to compensate for income losses from declining capture fisheries, disease-related mortalities have become a major challenge, causing significant economic losses for farmers. Due to inadequate screening of pathogens causing these mortalities, farmers have been left to rely on guesswork, frequently using treatments that are either ineffective or inappropriate. Therefore, this study aimed to identify and characterize the etiological agents responsible for infectious diseases in cage-farmed fish in Lake Victoria, Tanzania. Fish samples were collected from six districts in the Lake Victoria Basin (LVB), which are known for their extensive cage fish farming operations. A total of 81 samples of blood, liver, and kidney tissues were collected from morbid Nile tilapia and enriched in Buffered Peptone Water then cultured on various agar media to isolate pathogens. Gram staining and a series of biochemical tests were used to identify the bacterial isolates. Similarly genomic DNA were extracted from each isolate and fragments of the 16S rRNA gene amplified and sequenced. Three pathogenic bacteria namely *Citrobacter freundii, Pseudomonas aeruginosa,* and *Streptococcus agalactiae* were identified isolated and identified. Pathogenicity trials demonstrated that *P. aeruginosa* exhibited the highest mortality rate (86.7%), followed by *C. freundii* (66.7%) and *S. agalactiae* (40%). Clinical and post-mortem findings from the trials showed symptoms consistent with hemorrhagic septicemia and septicemia. Phylogenetic analysis grouped sequences of each pathogen into a single cluster, regardless of their geographical origins, suggesting a common source and subsequent dispersal to various locations. The study calls for harmonized efforts to enhance disease control strategies and reduce the impact on aquaculture operations, as farmers likely deal with the same bacterial strains.

## 1.0 Introduction

Fish diseases pose a serious constrain to the expansion and development of sustainable aquaculture (1). In global aquaculture, the emergence of previously uncharacterized pathogens linked to novel and unknown diseases has become an emerging trend (2). These pathogens typically spread rapidly, often transcending national boundaries, and cause major production losses approximately every 3–5years (2,3). The management efforts on the health of aquatic organisms have significantly increased during the last three decades (2). However, such efforts have not kept pace with the rapid growth of the aquaculture sector (4). Many of the most serious infectious disease agents affecting cultured species in aquaculture are bacteria (2). Bacteria rarely act as primary pathogens and are more commonly opportunistic pathogens in already immunocompromised hosts (5). Nevertheless, bacteria can cause significant losses in fish farming (2).

Cage fish farming was introduced in Lake Victoria (LV) in 2005, and since then, the region has seen a rapid increase in the number of cages, with over 8,024 now installed in the LVB (6). This method has become more intensive, with stocking densities reaching up to 150 kg/m³, compared to traditional pond systems and other aquaculture methods in the region, which yield between 0.2 and 100 kg/m³ (7). However, this expansion has been linked to significant losses from fish disease outbreaks, leading to fish morbidity and mortality. Morbidity results into poor growth, reduced market price of the produce, and high costs for treatment and preventive measures (7). According to the (8), global revenue losses attributed to fish diseases were estimated at 6 billion USD in the past ten years. In developing countries, over 50% of fish production is lost due to diseases, with China, a leading aquaculture producer, reporting 15% loss of total fish production due to diseases (5,9).

In Tanzania, less attention has been given to fish diseases in cage-farmed fish. Often, bruises, abrasions, and hemorrhages are incorrectly attributed to only overstocking rather than microbial infections. As a result, the use of salt (NaCl) treatments has become a common practice (10). A few studies focusing on bacteria are limited to the identification of pathogens at the genus level (11). This has left the status of fish bacterial pathogens and the associated diseases in cage fish farms poorly understood, increasing the likelihood that many fish diseases remain undiagnosed and continue to affect production. This knowledge gap leads to guesswork, with farmers often employing treatments that are incorrect or insufficient, thereby failing to address the actual causes of the diseases. (12). The delayed detection of diseases further compounds the issue, hindering the implementation of effective control measures (13). This deficiency accelerates the occurrence and spread of fish diseases within the cage fish farms of the Lake Victoria Basin (LVB) (14), leading to high mortality and discouraging farmers due to the increased economic loss. This study aimed to identify and characterise the different bacterial pathogens causing infectious diseases in cage-farmed fish in the Lake Victoria, Tanzania.

## 2.0 Materials and methods

### 2.1 Study area

The study was conducted in the Lake Victoria basin, Tanzania and it involved collection of infected fish samples from six districts along the shores of Lake Victoria in Tanzania: Nyamagana, Sengerema, Busega, Rorya Musoma MC, Musoma DC (Fig. 1). These districts were selected because they host a significant number of cage fish farming operations (15). The selected district host over 900 fish cages (16). The growth of cage fish farming in these districts has led to a substantial increase in fish production, contributing to the local economy and livelihoods (6). In response to this expansion, biosecurity measures are minimally employed through Environmental Impact Assessments (EIAs) conducted by the Tanzania Fisheries Research Institute (TAFIRI) and the National Environment Management Council (NEMC) to evaluate and mitigate the potential impacts of farming operations to safeguard both the environment and fish health (17). These necessitate formulation of basic biosecurity guidelines to be operationalized at cage level. Additionally, strategic areas for cage fish farming have been identified, and regulatory processes streamlined to support sustainable aquaculture practices.

**Figure1.**
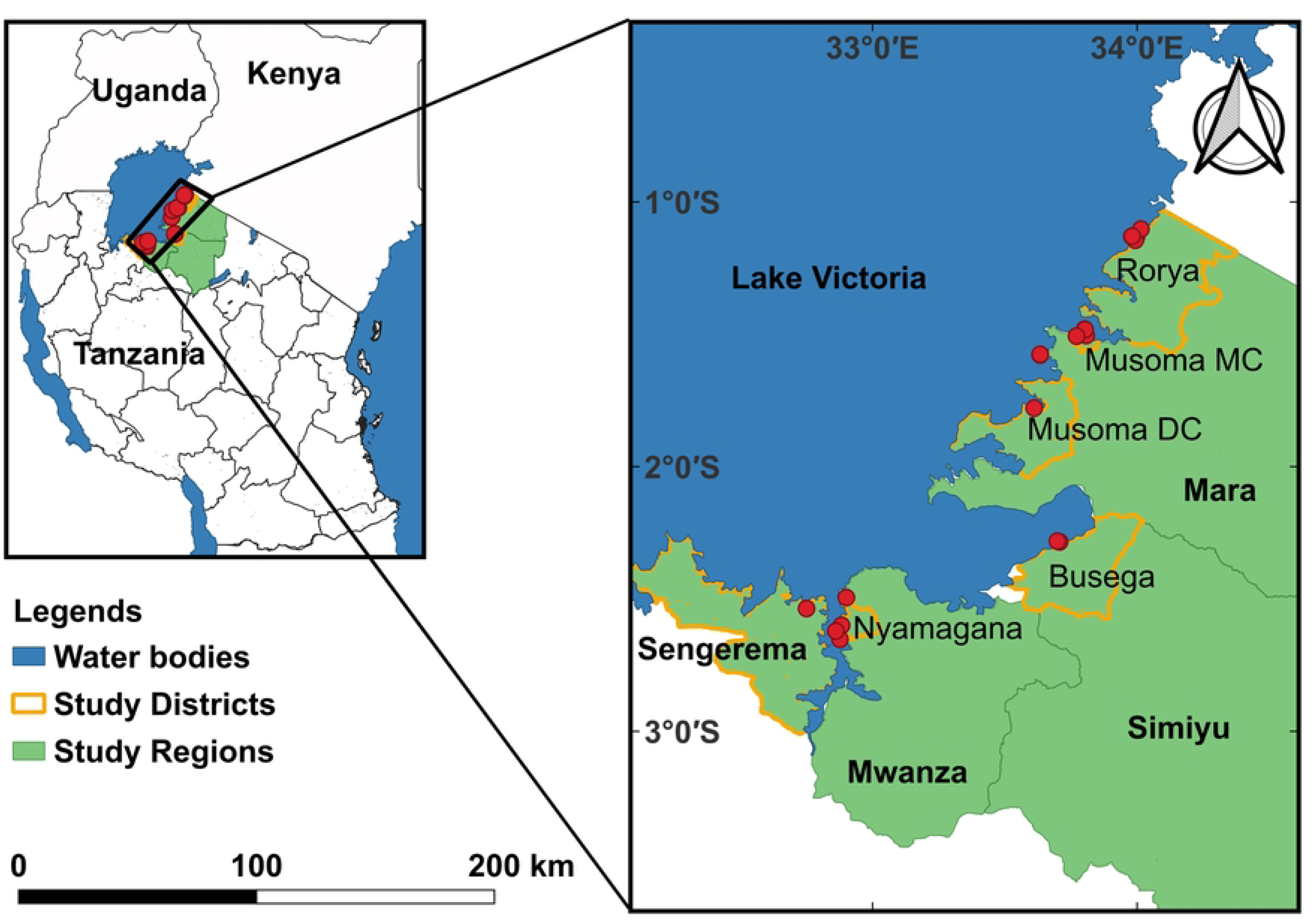
Map of the study area showing locations where bacterial samples were collected from infected fish. The map was created using QGIS software version 3.28 and shapefiles from the Database of Global Administrative Areas (GADM) (https://gadm.org/maps/TZA.html, accessed on 27th February 2025).

### 2.2 Ethical statement

The handling of fish and all the experimental protocols were carried out according to the guidelines of the Animal Ethics, established by the Sokoine University of Agriculture Research and Ethical Committee. To alleviate suffering during sampling fish were initially treated with MS-222 (Tricaine Methanesulfonate). The permit to conduct all experimental procedures and sampling was granted by Tanzania Commission for Science and Technology (COSTECH) while Tanzania Ministry of Regional Administration and Local Government granted authorisation for sample collection. Moreover, SUA approved and provided the permit to conduct the research under permit number SUA/DPRTC/R/126/CONAS/15/2023/14.

### 2.3 Fish sampling in the LVB cage fish farming systems

A total of eighty-one samples of Nile tilapia were collected from fish cages in the selected districts (Fig. 1). Priority was given to fish displaying clinical signs such as ulcerations, haemorrhage, bulging of the eyes, opaque eyes, eye loss, and scale loss according to (18). Samples of blood, liver, and kidney were taken for bacteria isolation according to (19). To preserve and maintain the viability of microorganisms during sample collection, Stuart Transport Medium was utilized as applied by (20). Cooler box was employed for insulation and temperature control to ensure that samples and transport media remain within the required temperature range (10 −15°C) during transportation. After sample collection, the samples were transported to Sokoine University of Agriculture for further analysis.

### 2.4 Isolation and identification of bacterial pathogens

Samples collected from fish with clinical signs were directly placed onto Buffered peptone water and incubated at 37 °C for 24 hours for enrichment. Then a loopful of incubated samples was taken and inoculated on Blood agar, *Pseudomonas* Lab agar, Mac Conkey agar, and Tryptic soy agar from Oxoid ltd media, UK and incubated at 37 °C for 24 hours, for isolation and identification of bacteria according to (10,18). All suspected colonies were purified by subculture for phenotypic and biochemical characteristics according to (21). All isolates were identified morphologically using Gram’s stain and biochemically using various biochemical tests, including enzyme tests such as catalase test and series of profiles of sugars such as Sulphide, indole, and motility test (SIM test), and Triple sugar iron (TSI) (10,18).

### 2.5 Molecular characterization of the isolated bacteria

The genomic DNA of the isolated Gram-negative bacteria was isolated using FlaPure Bacteria Genomic DNA Extraction Kit (Genesand Biotech Co., Ltd, Bei jing, China) following the manufacturer’s instructions. For the isolated Gram positive, the genomic DNA was isolated using the thermal extraction method in accordance to (22). A 1.0 mL of the Tryptic broth culture was pelleted, washed, and resuspended by vortexing in nuclease free water (Sourced from Inqaba biotech, Hatfield, South Africa), placed in a water bath at 95 °C for 5 min and immediately transferred to the ice for 5 min. This procedure was repeated, and the suspension was centrifuged at 10,000 rpm for 10 min (23). The quality of DNA extracts was checked on a 1 % agarose gel prior to subsequent analysis.

This was followed by amplification of the fragments of 16S rRNA (about 800 base pairs) in a T100 thermocycler using the universal primers 27[F:5’-AGAGTTTGATCATGGCTCAG-3’] and 1492 [R: 5’-TACGGYTACCTTGTTACGACTT-3’] (11). PCR was performed in a total volume of 35µL containing 17.5µL master mix, 1.75µL BSA, and 0.3 µM of each primer (23). The amplification was done as follows; Initial denaturation steps at 95 °C for 3 min and followed by 35 cycles of denaturation at 95 °C for the 30 s, annealing at 58 °C for 30 s and extension at 72 °C for 1 min followed by terminal extension at 72 °C for 3 min (10,23). The agarose gel (1.5%) stained with safe view was used to analyse PCR products by electrophoresis. Subsequently, successful PCR amplifications were sequenced using Sanger’s Dideoxy sequencing technology, (Microgen lab Amsterdam, Netherlands). The obtained 16S rRNA sequences were edited using the ClustalW algorithm as implemented in MEGA ver.11 (24). Using the Basic Local Alignment Search Tool (BLAST), each edited 16S rRNA sequence were compared to 16S rRNA barcode records published in the GenBank nucleotide database to confirm the taxonomic identity of each bacterium. FaBox DNA Collapser was used to collapse these sequences into unique haplotypes. Subsequently, the phylogenetic tree was constructed using BEAST ver 2.5 (25), to assess the evolutionary relationships among bacteria species. The analysis employed a relaxed uncorrelated log-normal molecular clock and a general time-reversible evolutionary model, running for 10 million generations. The phylogenetic tree was annotated using TreeAnnotator ver 1.10 (26) and visualized using FigTree ver 1.4 (27).

### 2.6 Pathogenicity test

#### 2.6.1 Acclimation period

A total of 200 healthy Nile tilapia weighing 50-200 g with no history of previous infections were collected randomly from Blue Economy Research Center, Sokoine University of Agriculture, Tanzania, and left acclimated in a clean round concrete tank of 3m^3^ holding capacity for seven days prior to pathogenicity experiments. Tanks were filled with underground well water with an average salinity of 0.3±0.1 g L^−1^. Dissolved oxygen was monitored at 5±1 mg L^−1^, while the water temperature was maintained at 27±0.52 °C according to (18). Tank pH was regulated at 7.5. The fish were fed two times daily (09:00 and 16:00 h) until visual satiety on a commercial pellet of 35% crude protein (Skretting, Alexandria, Egypt) according to (18). The organic wastes and other debris were siphoned and 30% of the water was replaced daily to reduce the toxicity of ammonia. Fish that showed normal reflexes with no apparent lesions were selected pathogenicity assessment.

#### 2.6.2 Pathogenicity assessment

180 of acclimated Nile tilapia were equally distributed into twelve concrete tanks; each tank stocked with 15 fish. The fish of the first group (C) were not assigned to any treatment and served as a control, while the fish of the other groups (T1, T2 and T3) were injected with 0.5 mls of the overnight culture of virulent *C. freundii*, *P. aeruginosa*, and *S. agalactiae* respectively at a concentration of 3×10^7^ cells mL^−1^ (Fig. 2), cultured on tryptic soy broth (Oxoid) at 37 °C for 24 h accordance to (18). To prevent unnecessary suffering, fish exhibiting erratic swimming, hanging motion, and aneroxia were removed immediately (15-30 minutes) from experiment. The pathological clinical signs and cumulative deaths were recorded daily among experimental groups for two weeks post-treatment. Sample from moribund and freshly dead fish were collected, examined immediately to verify the cause of death.

**Figure 2:**
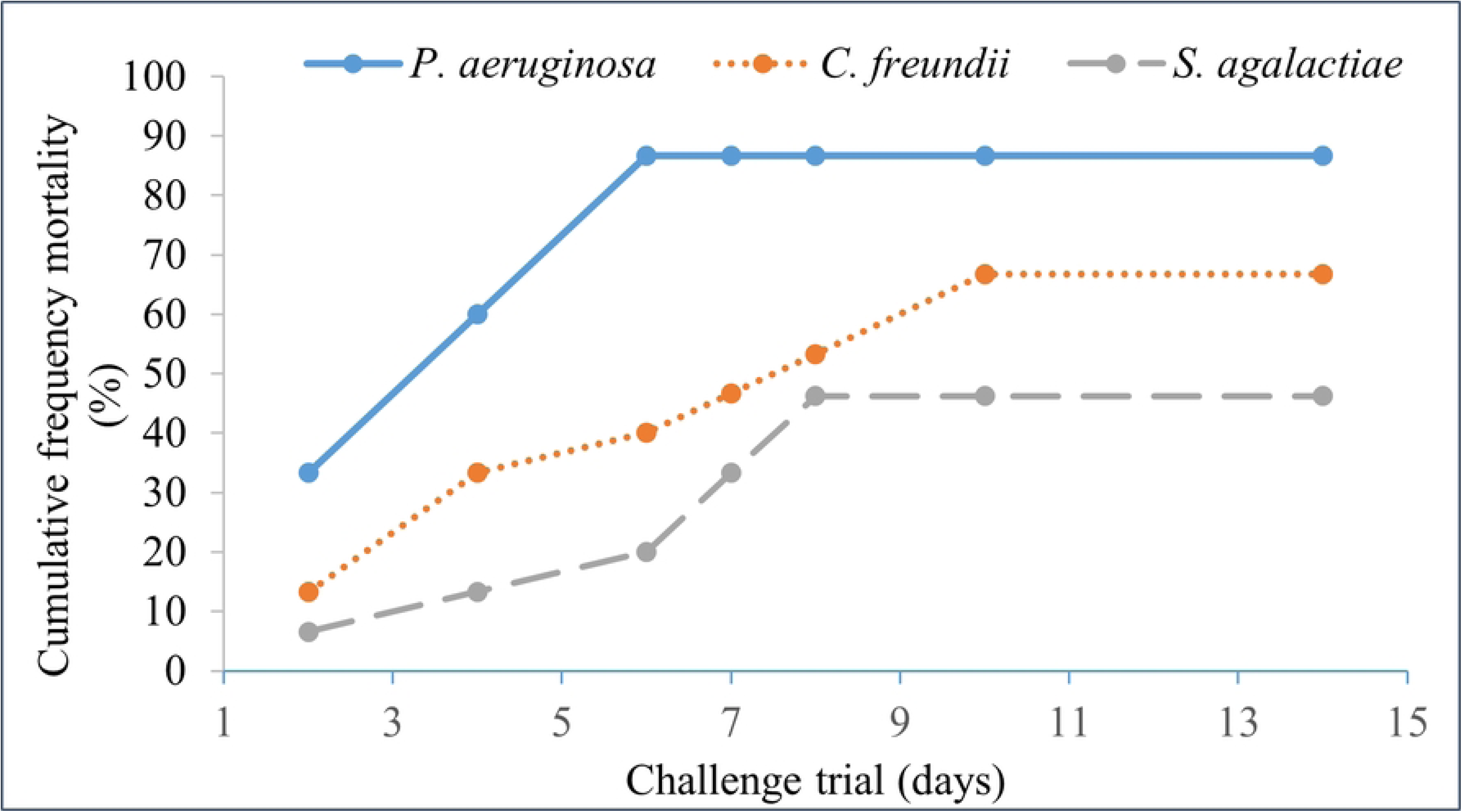
Experimental design for Pathogenicity test; Control (group not inoculated with any bacteria), T1 (Test group inoculated Intraperitonealy with *C. freundii*), T2 (Test group inoculated Intraperitonealy with *P. aeruginosa*) and T2 (Test group inoculated Intraperitonealy with *S. agalactiae*)

## 3.0 Results

### 3.1. Bacteriological assay

The bacteriological examination identified the primary bacteria affecting fish in the Nyamagana, Rorya, and Musoma MC districts as *Citrobacter* spp., *Pseudomonas* spp., and *Streptococcus spp.* Specifically, all examined samples from Sengerema tested positive for *Citrobacter spp.*; samples from Musoma DC revealed *Pseudomonas spp.* as the dominant bacteria; and samples from Busega district tested positive for both *Citrobacter spp.* and *Pseudomonas spp.* All retrieved isolates were motile, Gram-negative bacilli, arranged in double or short chains. The colonies reacted positively to catalase, oxidase, nitrate reduction, gelatin hydrolysis, citrate utilization, and mannitol fermentation, while they were negative for H_2_S production, urease, indole, methyl red, voges Proskauer and citrate utilization test except for samples were *Streptococcus* spp., isolated. The typical isolates of *Psedomonas* spp., displayed large, irregular colonies with a fruity odor and produced a yellowish-green fluorescent pigment on *Pseudomonas* Lab Agar after 24 hours at 37 °C (figure 3a). On *Pseudomonas* Lab Agar and MacConkey Agar, the bacteria formed flat, non-lactose-fermenting colonies with regular edges and an alligator skin-like appearance from the top view. The isolate of *Citrobacter* spp., on Tryptic Soy Agar showed yellowish, opaque, round, convex, smooth-edged, and semi-translucent colonies (figure 3b). The presumed *Streptococcus* spp were non-motile, Gram-postive cocci, arranged in chains; the colonies reacted negatively to catalase and SIM test that included Sulphide, indole, and motility test (figure 3c). Based on morphological and biochemical characteristics, all isolates were identified as *Citrobacter* spp., *Pseudomonas* spp*., and Streptococcus* spp.

**Figure 3:**
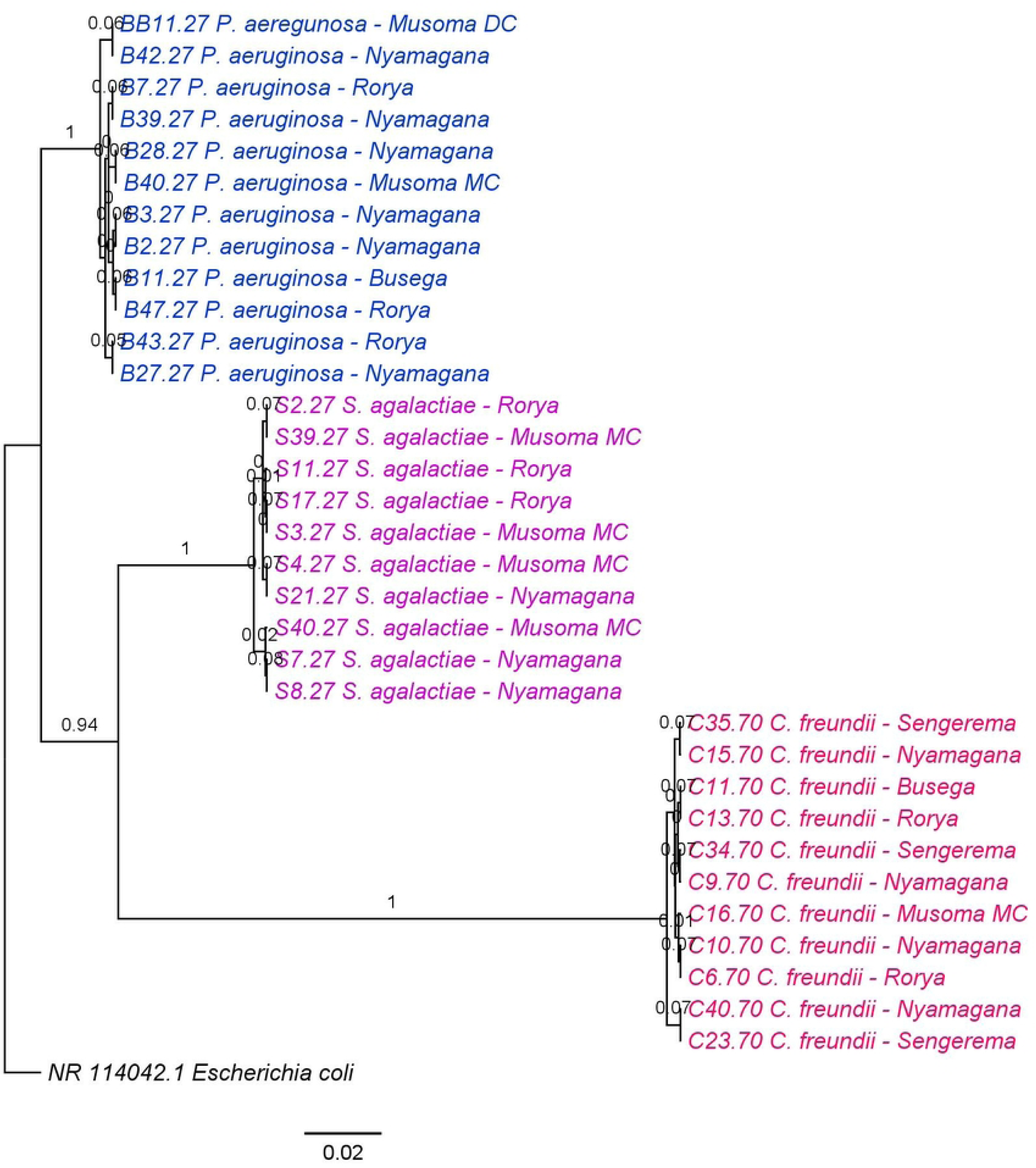
Colonies of isolated bacteria. **(a)** *Pseudomonas* spp with large irregular colonies produced a yellowish-green florescent pigment on Pseudomonas Lab agar. **(b)** *Citrobacter* spp., on Tryptic soy agar showed yellowish opaque, round, convex, smooth edged and semi-translucent colonies. **(c)** *Streptococcus* spp. isolates were non-motile, Gram-postive coccus, arranged in chains.

### 3.2 Barcoding and distribution of bacteria isolated from cage fish farms

Bacteriological examination revealed that (37%, n=35) of all positive samples were collected from Nyamagana district, followed by Rorya district (23%, n=35), Musoma MC (17%, n=35), Busega and Sengerema (9% each, n=35), and Musoma DC, which recorded the lowest percentage (6%, n=35). Regarding bacterial isolates, *C. freundii* accounted for 31% of all isolates. Its distribution varied across districts, with Nyamagana contributing (11%, n=11), Rorya (6%, n=11), Sengerema (9%, n=11), and Musoma DC, Busega, and Musoma MC each contributing (3%, n=11). *P. aeruginosa* was the most prevalent bacterial isolate, comprising 40%, (n=14) of all isolates, with contributions from Nyamagana (17%, n=14), Musoma MC (9%, n=14), Busega (6%, n=14), Musoma DC (6%, n=14), and (Rorya 3%, n=14). *S. agalactiae* represented 29% of all isolates, found in Nyamagana (9%, n=10), Rorya (9%, n=10), and Musoma MC (11%, n=10) (Table 1). The analysis of 43 16S rRNA sequences of bacteria revealed that 35 of these sequences were associated with three bacterial species: *C. freundii*, *P. aeruginosa*, and *S. agalactiae* (Table 2). The BLAST results showed percentage identities ranging from 98.29% to 100%, with expected values of 0 for all sequences. Maximum scores ranged from 811 to 1109, while query coverage was 100% for each blasted sequence.

**Table 1:** Distribution of bacteria isolated from cage fish farms in selected districts in Lake Victoria basin, Tanzania.

|  | Nyamagana | Sengerema | Musoma | Musoma | Rorya | Busega |
| --- | --- | --- | --- | --- | --- | --- |
|  |  |  | MC | DC |  |  |
| *S. agalactiae* (%) | 9 | 0 | 11 | 0 | 9 | 0 |
| *C. freundii* (%) | 11 | 9 | 3 | 0 | 6 | 3 |
| *P. aeruginosa* (%) | 17 | 0 | 9 | 6 | 3 | 6 |

**Table 2:** BLAST results obtained after comparing the 16S rRNA haplotypes sequences of isolated bacteria samples from the Lake Victoria cage fish farms, with those in the NCBI database. E-value, Expected value.

| Haplotype | Species | Base pair size | Maximum | Query | E- | % |
| --- | --- | --- | --- | --- | --- | --- |
| (H) |  |  | score | cover (%) | value | Identity |
| H1 | *P. aeruginosa* | 600 | 1109 | 100 | 0.0 | 100 |
| H2 | *S. agalactiae* | 600 | 1103 | 100 | 0.0 | 99.83 |
| H3 | *S. agalactiae* | 600 | 1109 | 100 | 0.0 | 100 |
| H4 | *C. freundii* | 600 | 811 | 100 | 0.0 | 98.29 |

### 3.3 Phylogenetic analysis

The phylogenetic tree of 16S rRNA gene clustered the analysed sequences together for each species of bacteria whose taxonomic identity is reliable (Figure 4), indicating accurate species identification through the amplified barcode. Similarly, bacteria of the same species clustered together irrespective of their geographical origin (Figure 4) demonstrating strong genetic conservation within bacterial species across sampling sites in cage fish farms within Lake Victoria. This finding suggests that geographical separation has minimal impact on the genetic structure of these bacterial populations.

**Figure 4.**
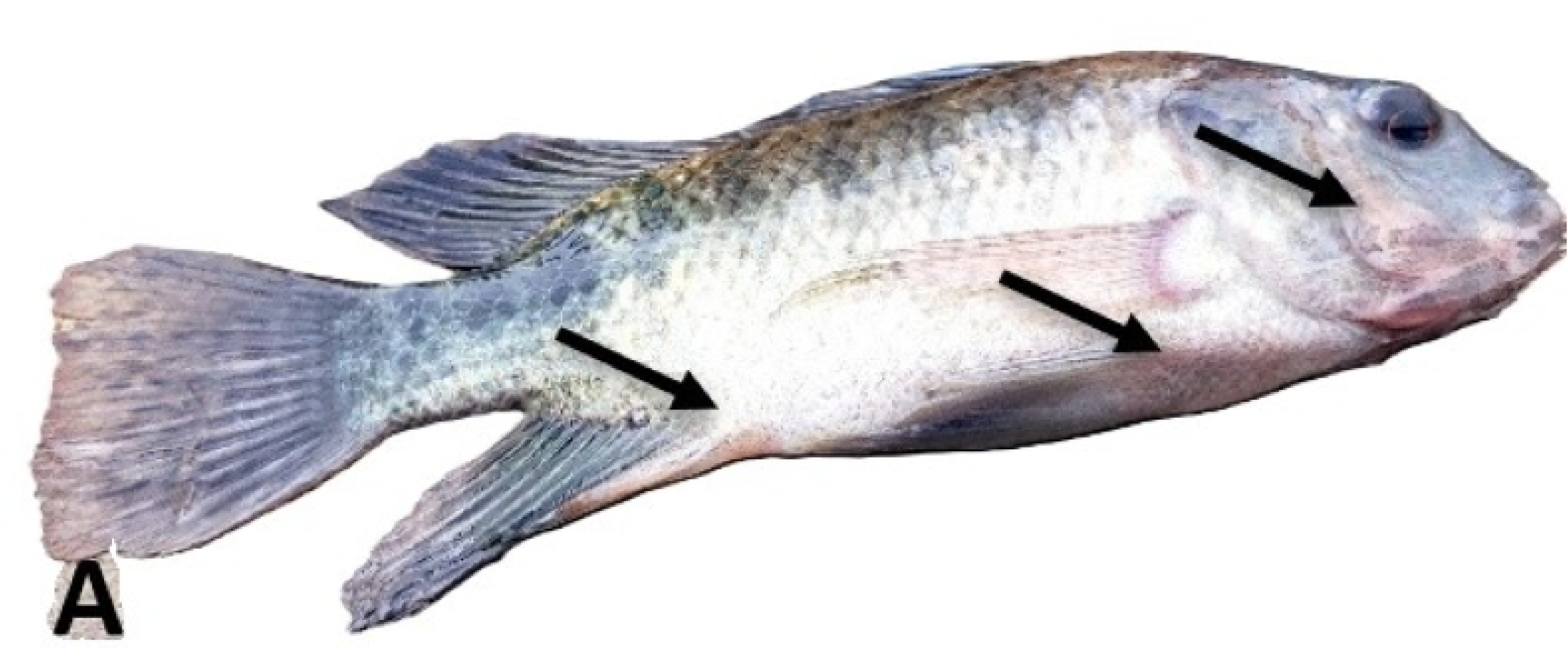
Phylogenetic tree showing the evolutionary relationships among 16S rRNA haplotypes of pathogenic bacteria isolated from Nile tilapia sampled from cage fish farms in the Lake Victoria.

### 3.4 Pathogenicity assessment

The clinical signs as well as fish morbidity and mortality were recorded in all experimental groups for two weeks post-challenge. The results demonstrated that the fish of the control group did not reveal any mortalities or pathological lesions, while those of the other groups displaying high mortalities and pathological lesions of hemorrhagic septicemia, similar to those reported in naturally infected fish. Fish exposed to *P. aeruginosa,* showed ragged fin, gill erosion, pectoral fin hemorrhage (Figure 6b)., white pectoral fin, redding of the fin that develop to white patches, detached scales and scattered hemorrhagic spots (Figure 6a). Those exposed to *S. agalactiae,* showed scale loss, eye hemorrhage, fin rot (Figure 8a), bulging of the eye, cloudy aye, aye loss, white color to the base of openings and fins, (Figure 8b) while those exposed to *C. freundii*, showed skin hemorrhage, fin rot, and skin ulcer which ended up into open sore (Figure 7). The maximum mortality rate (86.7%) was recorded in fish exposed to *P. aeruginosa*, followed by *C. freundii*, (66.7%) and the least was observed in *S. agalactiae* (40%). Indeed, Nile tilapia exposed to *P. aeruginosa*, produced higher mortalities in a shorter time compared to those exposed to *C. freundii*, and *S. agalactiae* (Figure 5). Approximately, 87.5% of fish exposed to *P. aeruginosa* died within six days post-inoculation, while those of exposed to *C. freundii*, and *S. agalactiae* showed delayed mortality of 67.5% and 40% respectively up to ten days post-inoculation. The postmortem findings revealed that the all-exposed fish displayed typical signs of septicemia manifested by congested liver, enlarged spleen, and accumulation of serous bloody fluid in the abdominal cavity. In terms of bacteriology, *C. freundii*, *P. aeruginosa*, and *S. agalactiae* ware successfully reisolated from skin ulcers and internal organs of dead and moribund fish, and the results confirmed that all isolates belonged to *C. freundii*, *P. aeruginosa*, and *S. agalactiae*.

**Figure 5:**
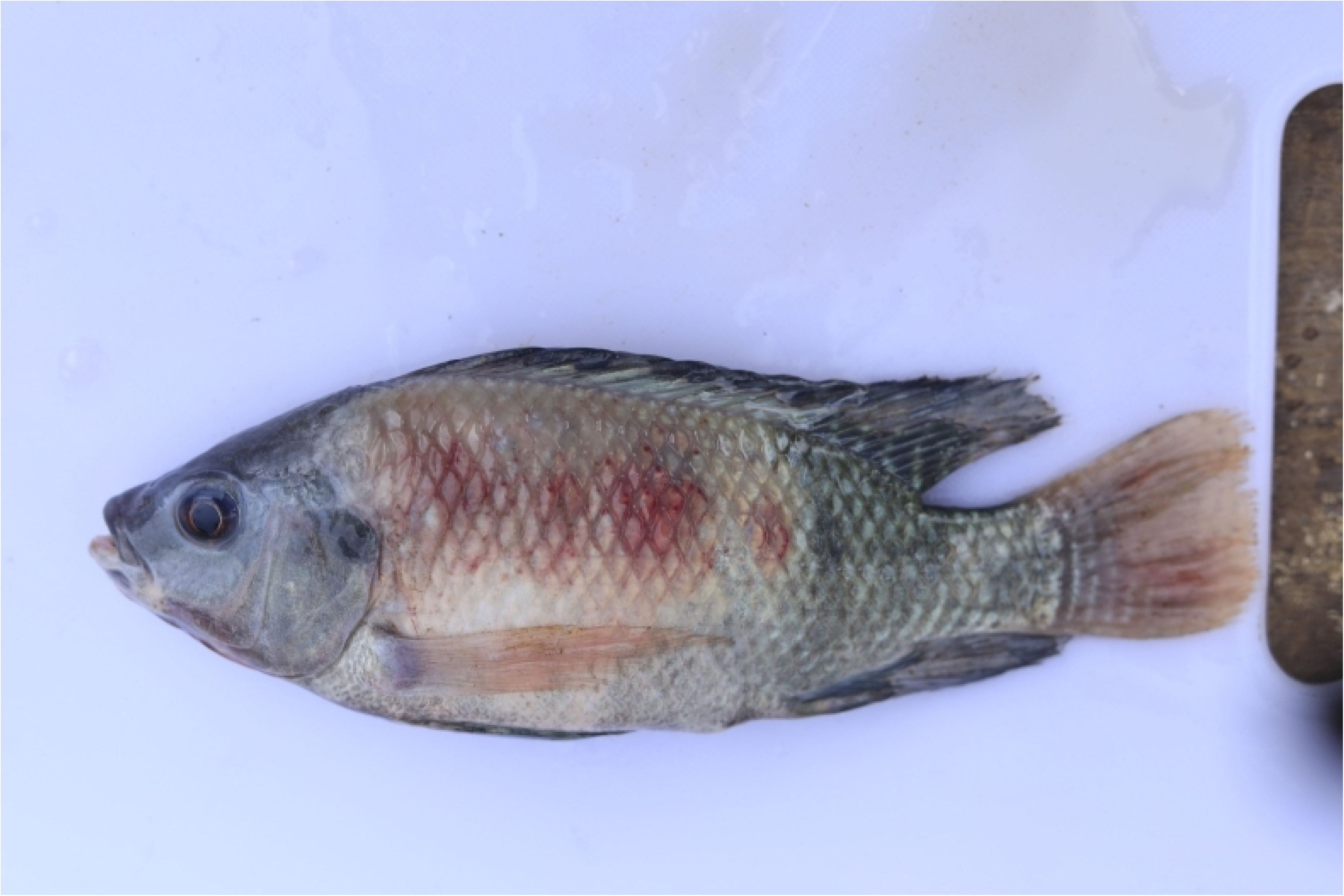
Cumulative mortality of Nile tilapia inoculated intraperitoneally with 0.5 ml of isolated bacteria at a concentration of 3×10^7^cfu ml^−1^

**Figure 6:**
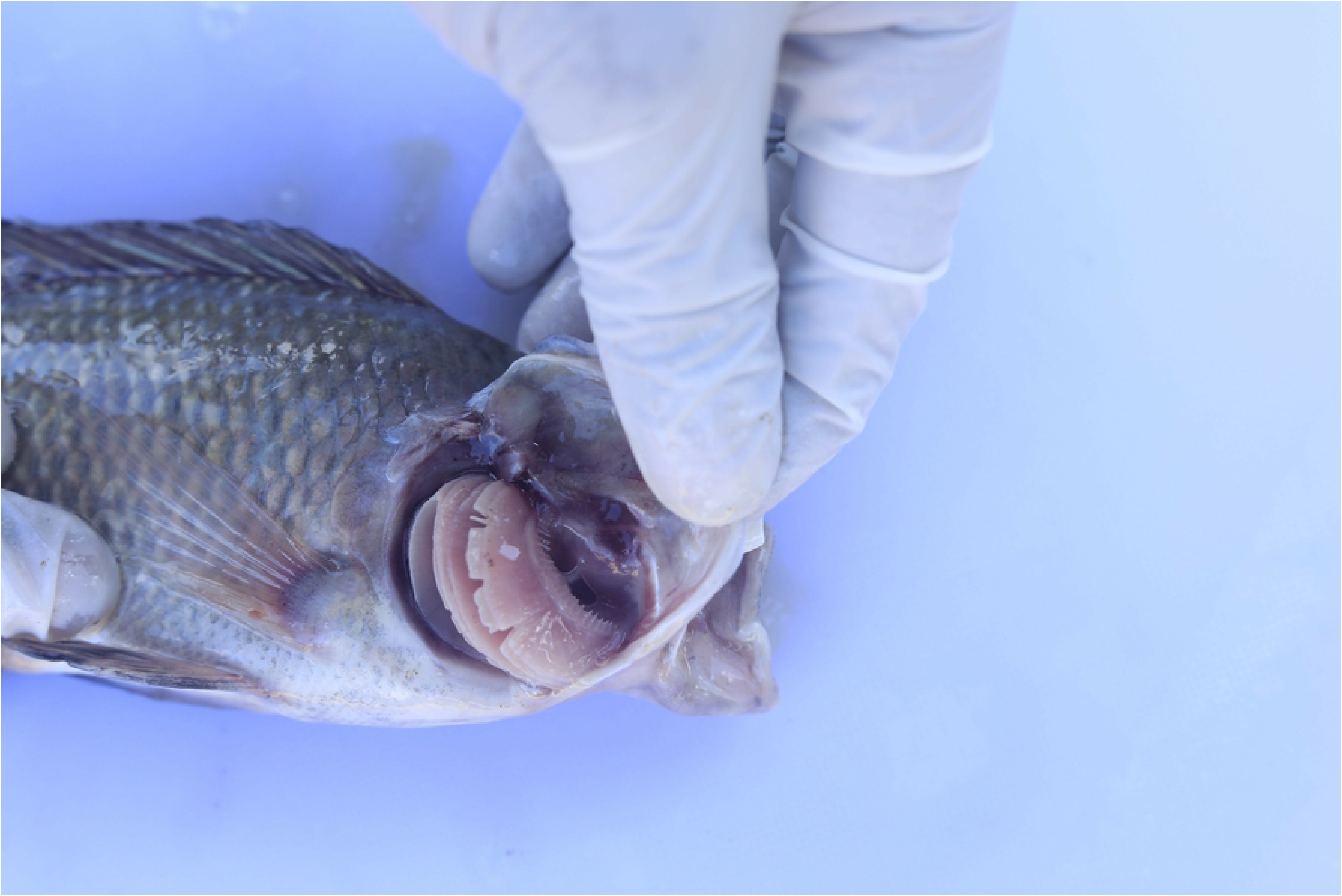
Clinical examination of fish exposed to *P. aeruginosa*, **(a)** Nile tilapia (*O. niloticus*) showing scattered hemorrhagic spots (black arrows) **(b)** Nile tilapia (*O. niloticus*) showing scattered hemorrhagic spots, detached scales, gill erosion and fins erosion.

**Figure 7:**
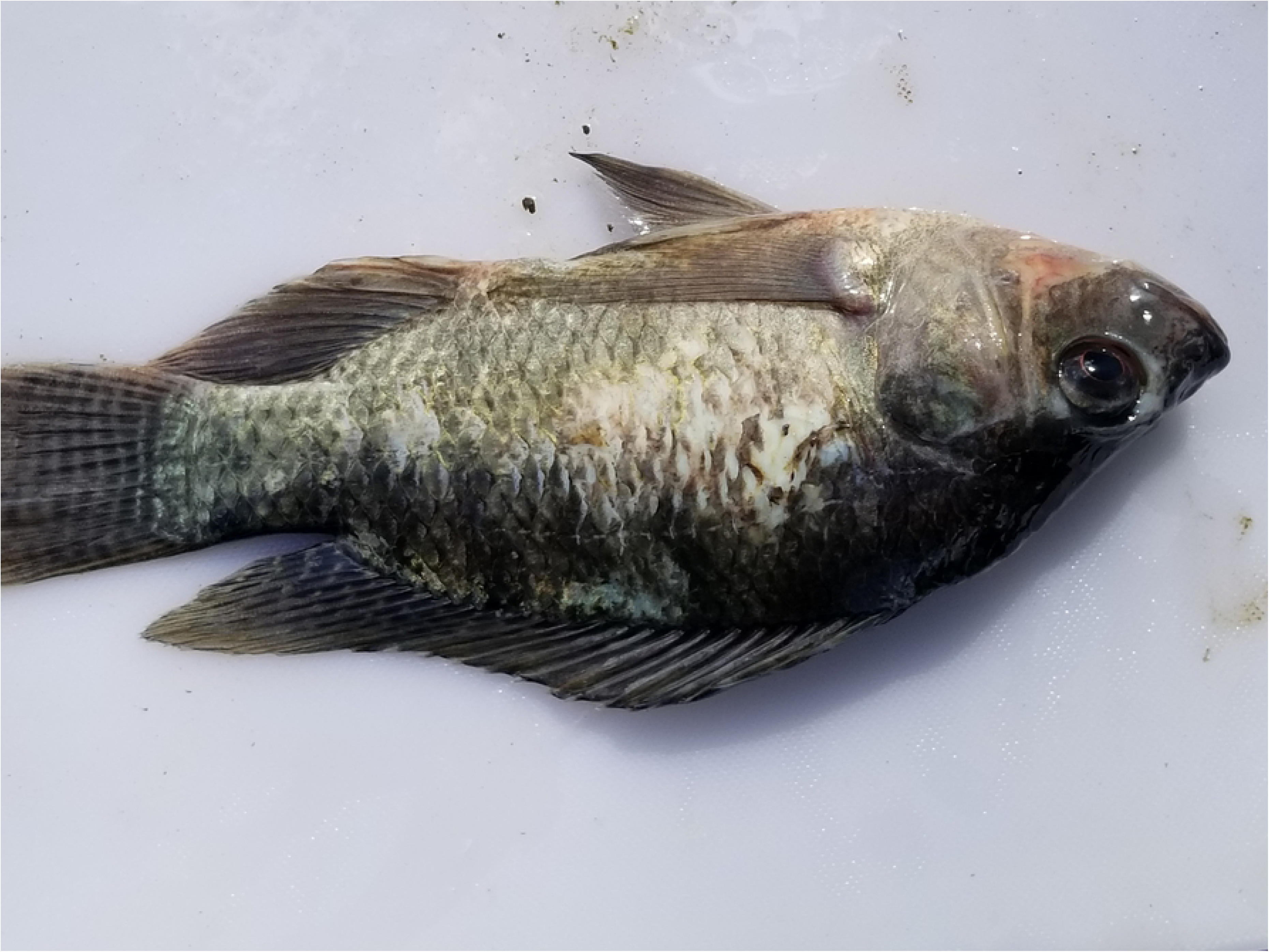
Clinical examination of fish exposed to *C. freundii* fish showing ragged fin hemorrhage, skin hemorrhage and fins erosion

**Figure 8:**
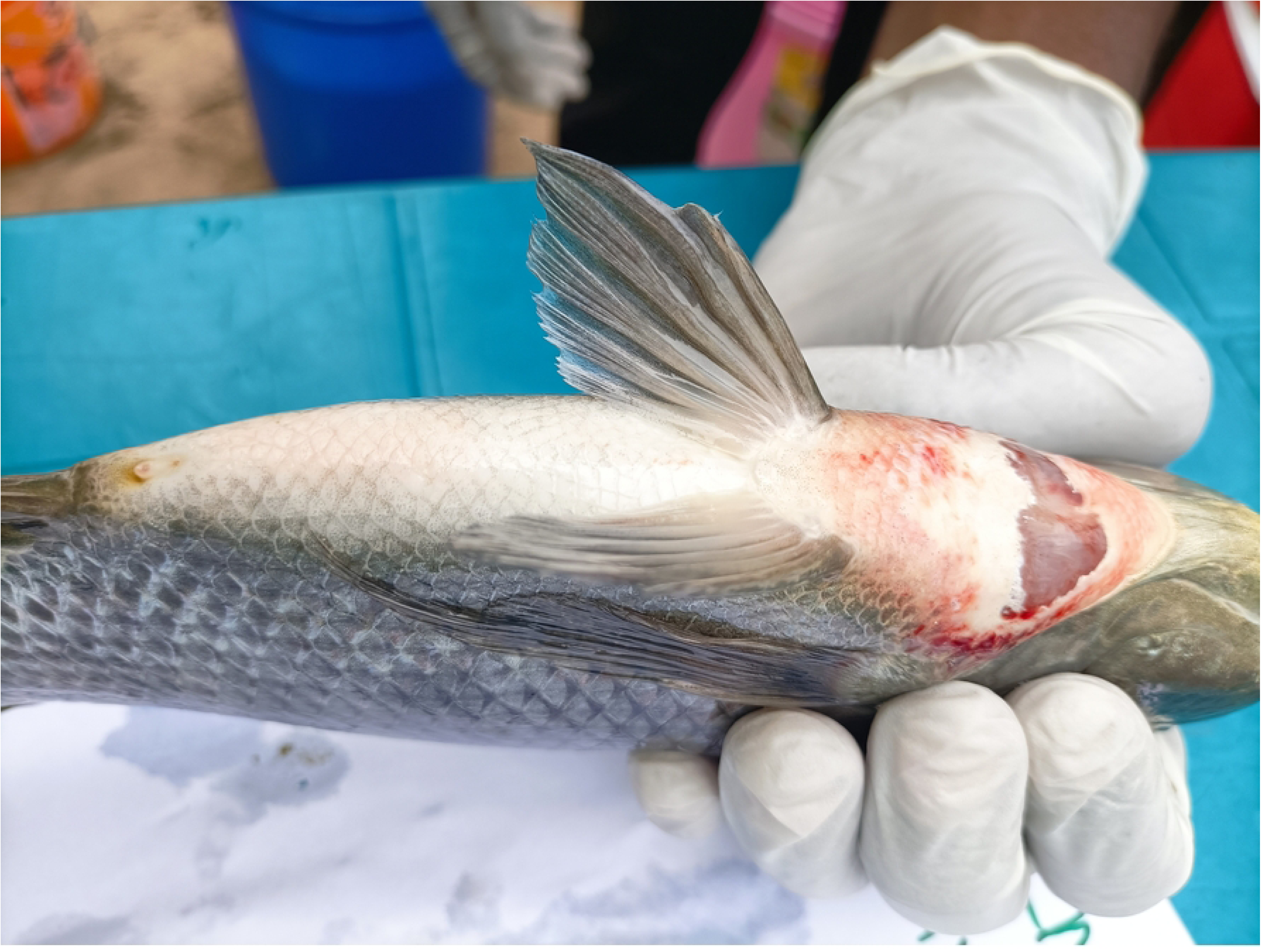
Clinical examination of fish exposed to *S. agalactiae.* **(a)** fish showing scale loss, ulcers and fins erosion. **(b)** Fish showing aye loss.

### 3.5 Clinical and postmortem findings of fish collected in cage fish farms

A total of 81 morbid fish were collected. The clinical inspection revealed that most of the naturally infected fish shared the same typical clinical signs, including hemorrhages on external body surfaces, mainly at the ventral aspect of the abdomen and around the vent. Others showed fins erosions, skin darkness, ulcers, aye loss, scale loss and detached scales (Fig. 9). Internally, the infected fish showed typical signs of hemorrhagic septicemia, pale liver, necrotic gills, and empty GIT.

**Figure: 9.**
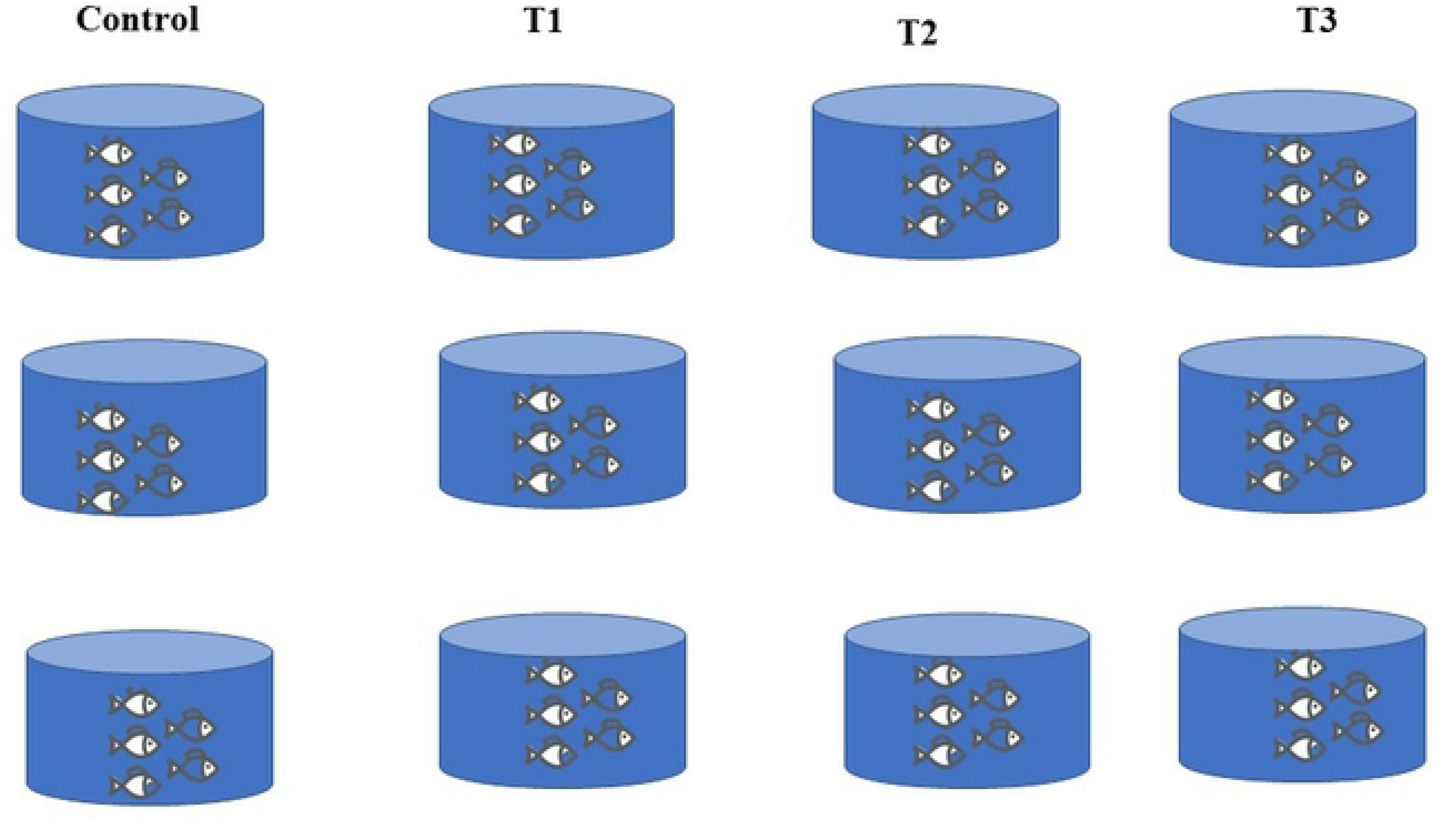
Clinical examinations of naturally infected fish Nile tilapia (*O. niloticus*) showing skin ulcer.

## 4.0 Discussion

The study identified *P. aeruginosa*, *C. freundii*, and *S. agalactiae* as the major bacterial species affecting fish in cage fish farms in the LVB. Notably, *P. aeruginosa* was predominantly isolated from Nyamagana, Rorya, Musoma MC, Musoma DC, and Busega, and accounted for 40% of the bacterial isolates, making it the most common pathogen. This high prevalence aligns with previous studies that reported *P. aeruginosa* as a major contributor to bacterial infections in fish farms, accounting for over 50% of such cases from tilapia and channel catfish cultured in Indonesia (28–30). The widespread presence of *P. aeruginosa* in the LVB can be attributed to its ubiquitous nature in aquatic environments, as well as similarities in production systems and fish species across the sampled districts (28,30). Symptoms observed in fish exposed to *P. aeruginosa* included ragged fins, gill erosion, hemorrhages, and white patches, consistent with findings from previous studies (2,31). Similarly, the pathogen resulted in the highest mortality rate (86.7%) and rapid onset of disease compared to other isolated pathogens. Moreover, *Pseudomonas* spp., particularly *P. aeruginosa* and *P. fluorescens*, are known to be significant pathogens in both marine and freshwater aquaculture systems (11,32). The association of *P. aeruginosa* with poor water quality in fish farms, further highlights the need for improved management practices (33).

*Citrobacter freundii* accounted for 31% of the bacterial isolates and was isolated in samples collected from all districts except Musoma DC, indicating their widespread prevalence. Infections caused by *C. freundii* are known to result in severe conditions such as hemorrhagic septicemia, edema, epizootic ulcerative syndrome (EUS), hemorrhagic enteritis, and red body disease, affecting a variety of finfish species, including common carp, goldfish, eel, catfish, and tilapia (28). Although *C. freundii* caused significant fish mortality following the pathogenicity assessment (66.7%), the progression of the disease was somewhat slower compared to *P. aeruginosa*. Symptoms observed included skin hemorrhages, fin rot, and skin ulcers, consistent with other studies (34). These findings emphasize the importance of addressing *C. freundii* infections to mitigate their impact on fish health and aquaculture productivity.

*Streptococcus agalactiae* was isolated in 29% of the samples and was primarily limited to Musoma MC, Rorya, and Nyamagana. Symptoms associated with *S. agalactiae* infections included scale loss, eye hemorrhage, and fin rot, consistent with other findings which evaluated the clinical and histopathological lesions associated with the bacteria (35–37). Despite a lower mortality rate of 40%, over 60% of fish infected with *S. agalactiae* recovered, though often with permanent eye damage. This impairment can negatively affect feeding responses, leading to reduced growth and increased feed wastage. Accumulated uneaten feed can further deteriorates water quality, potentially causing economic losses for cage farmers and exacerbating environmental issues (37,38). Although *P. aeruginosa* remains the most prevalent and problematic pathogen, *C. freundii* and *S. agalactiae* also pose significant risks to fish health in cage fish farming in the study area.

The clustering of *S. agalactiae*, *P. aeruginosa*, and *C. freundii* isolates into distinct phylogenetic groups across different districts highlights the potential for a shared source of infection among fish farms. These results align with findings from previous studies emphasizing the role of interconnected aquaculture practices and environmental factors in disease transmission. For instance, the genome-based analysis of multidrug-resistant *E. coli* in Lake Victoria (39) illustrated how shared water systems can serve as reservoirs for pathogens, facilitating their dissemination across districts. Similarly, studies on *S. agalactiae*, and *P. aeruginosa* in fish farms in Brazil (21) and Indonesian (40) reveal the genetic uniformity of strains within localized outbreaks, suggesting common sources of infection, such as contaminated feed, equipment, or water. This is consistent with our findings, where clustering patterns imply that a single strain of each bacterial species is circulating across multiple districts. Moreover, the molecular identification of pathogens in Nile tilapia and the comparison of *S. agalactiae*, *C. freundii* and *P. aeruginosa* isolates from different hosts underline the adaptability and resilience of these bacteria within aquaculture systems (41,42). The presence of identical strains across districts in our study further supports the hypothesis that shared resources such as water systems and farming practices play a significant role in pathogen transmission (39,40). Implementation of unified disease management measures across districts, as farmers are likely dealing with the same bacterial strains and coordinated efforts could enhance the effectiveness of disease control strategies and reduce the impact on cage fish farming operations in the study area.

## 5.0 Conclusion

This study revealed the presence of three pathogenic bacteria in cage fish farming systems in Lake Victoria, with *P. aeruginosa* being the most prevalent and problematic pathogen, affecting fish across all sampled districts except Sengerema districts. *Citrobacter freundii* and *S. agalactiae* also pose significant threats, with *P. aeruginosa* causing the highest mortality rates and *S. agalactiae* impairing fish recovery and growth. These pathogens exhibit distinct clinical manifestations, yet collectively contribute to substantial health challenges in the study area. Phylogenetic analysis showed that the clustering of *S. agalactiae*, *P. aeruginosa*, and *C. freundii* isolates into distinct phylogenetic groups across different districts highlights the potential for a shared source of infection among fish farms. These findings emphasize the urgent need for harmonized management practices in the LVB and biosecurity measures in cage fish farming to mitigate the impact of bacterial infections. Strengthening biosecurity protocols, improving water quality, and implementing robust disease monitoring systems are also crucial in reducing pathogen prevalence and their harmful effects. Similarly, routine screening for pathogenic diseases in cage fish farms and hatcheries should be enforced to ensure biosecurity compliance and disease control. Further research should be conducted to track the root cause of bacterial diseases in the LVB by investigating environmental reservoirs, transmission routes, and the role of wild fish species in pathogen dissemination. Addressing these issues will not only improve fish health but also minimize the economic losses faced by farmers in cage fish farming systems in the Lake Victoria.

## Acknowledgment

We would like to express our gratitude to cage fish farmers in the LVB for their support during fieldwork. Furthermore, we extend our appreciation to the SUA and the Tanzanian Ministry of Regional Administration and Local Government for issuing the necessary permits. Furthermore, the authors are thankful to Eric Rwiza Oswad and Mikidadi Rashidi Abdallah from the college of veterinary and Biomedical sciences for their support during laboratory analysis of sample. Colleagues from the Department of Biosciences, SUA are also gratefully thanked for their support during molecular analysis of the samples.

---

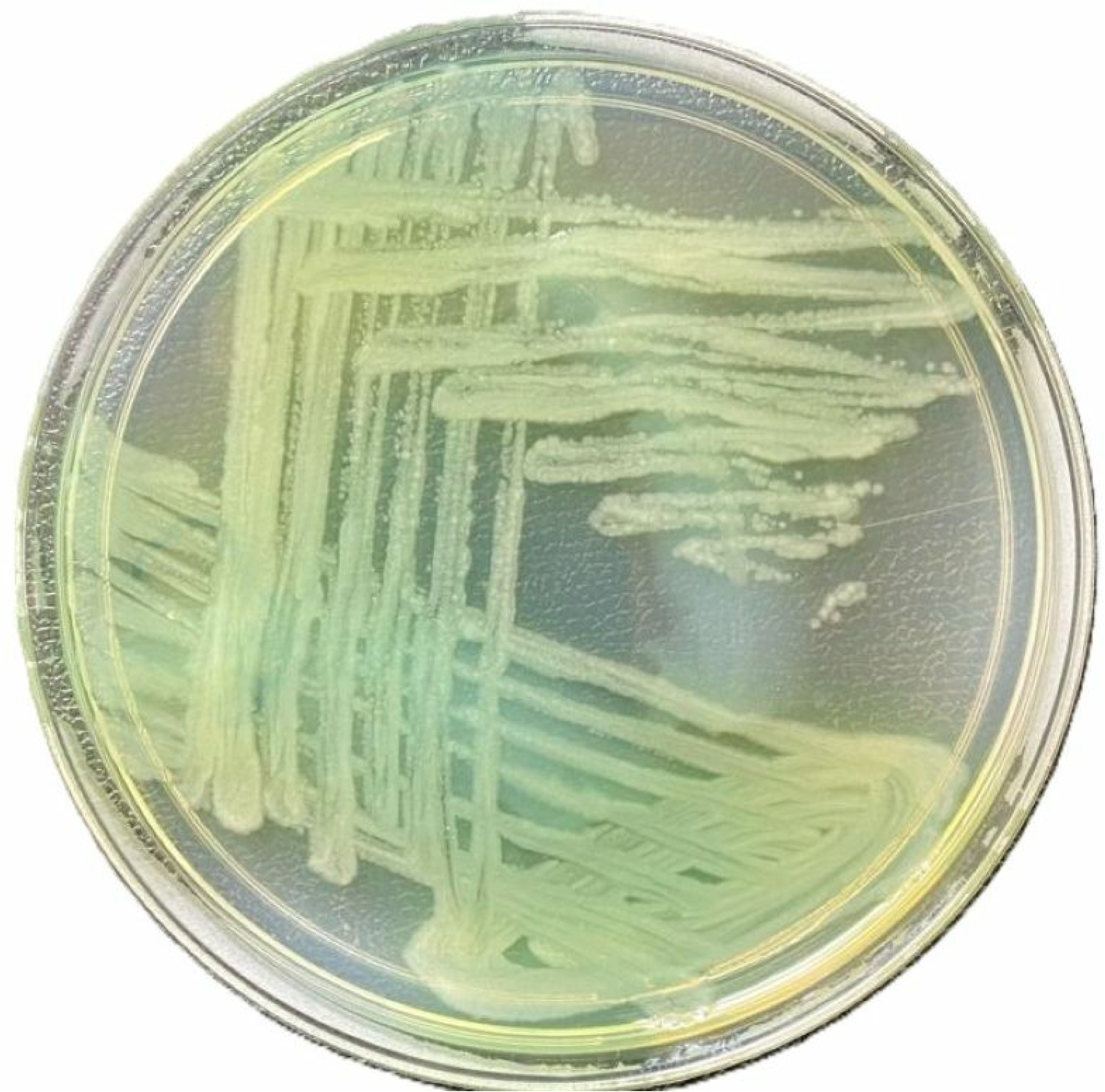

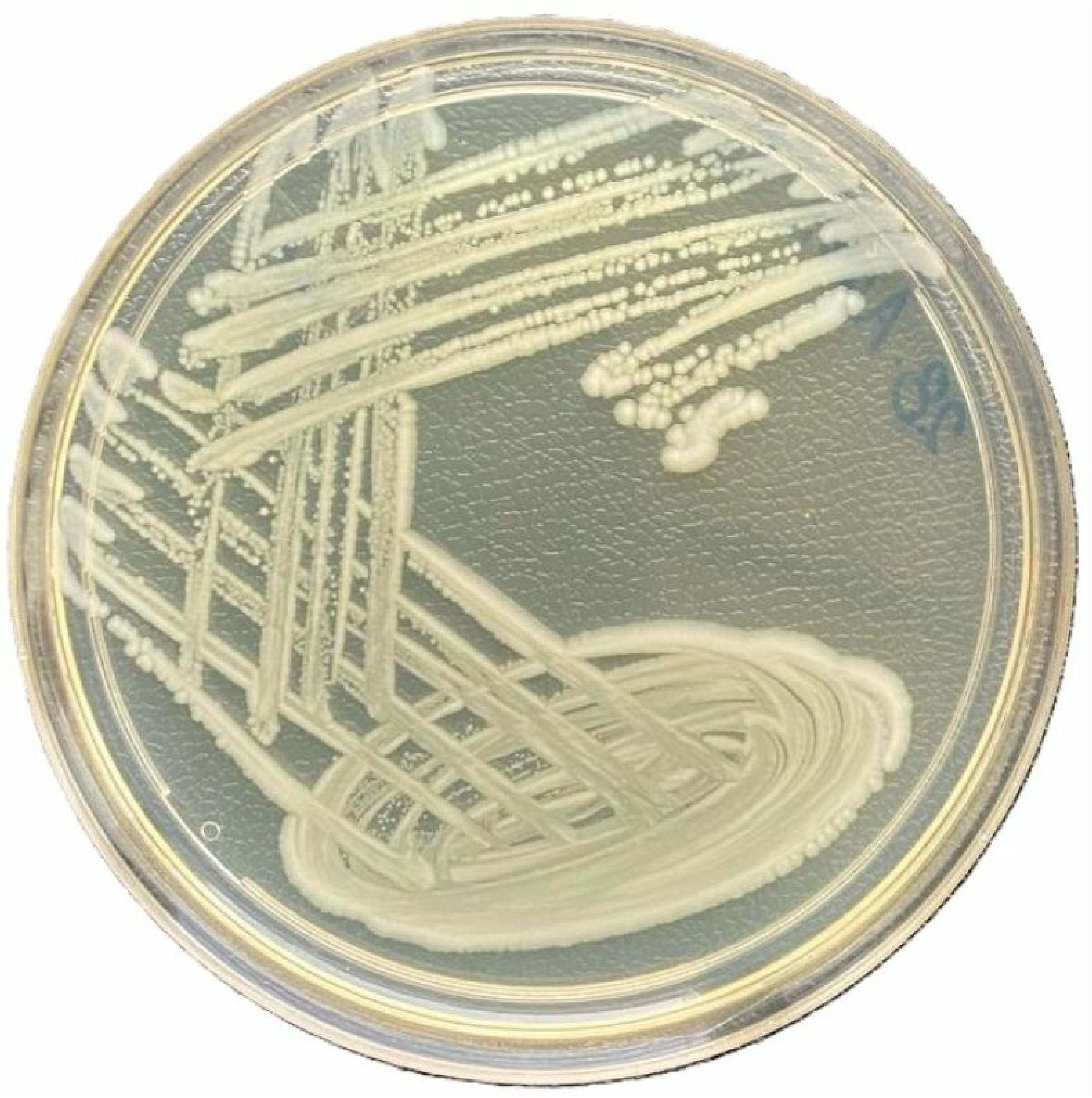

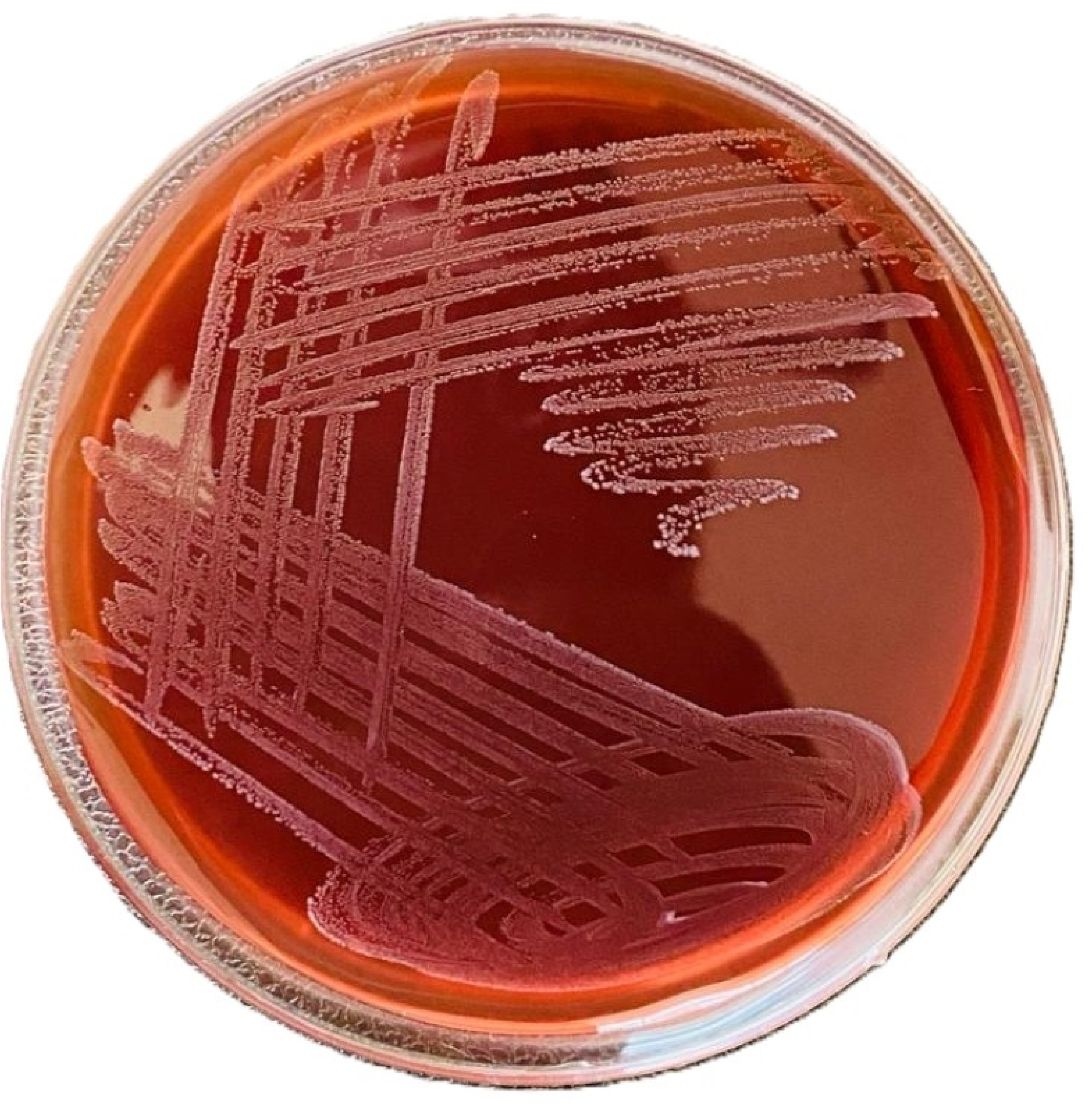

## Notes

### Competing Interest Statement

The authors have declared no competing interest.

